# Spontaneous emergence of music detectors in a deep neural network

**DOI:** 10.1101/2021.10.27.466049

**Authors:** Gwangsu Kim, Dong-Kyum Kim, Hawoong Jeong

## Abstract

Music exists in almost every society, has universal acoustic features, and is processed by distinct neural circuits in humans even with no experience of musical training. These characteristics suggest an innateness of the sense of music in our brain, but it is unclear how this innateness emerges and what functions it has. Here, using an artificial deep neural network that models the auditory information processing of the brain, we show that units tuned to music can spontaneously emerge by learning natural sound detection, even without learning music. By simulating the responses of network units to 35,487 natural sounds in 527 categories, we found that various subclasses of music are strongly clustered in the embedding space, and that this clustering arises from the music-selective response of the network units. The music-selective units encoded the temporal structure of music in multiple timescales, following the population-level response characteristics observed in the brain. We confirmed that the process of generalization is critical for the emergence of music-selectivity and that music-selectivity can work as a functional basis for the generalization of natural sound, thereby elucidating its origin. These findings suggest that our sense of music can be innate, universally shaped by evolutionary adaptation to process natural sound.

**One-sentence summary:** Music-selectivity can arise spontaneously in deep neural networks trained for natural sound detection without learning music.

## MAIN

Music is a cultural universal of all human beings, having common elements found worldwide^1,2^, but it is unclear how such universality arises. As the perception and production of music stem from the ability of our brain to process the information about musical elements^3–7^, the universality question is closely related to how neural circuits for processing music develop, and how universals arise during the developmental process regardless of the diversification of neural circuits derived by the spectacular variety of sensory inputs from different cultures and societies.

In our brain, music is processed by music-selective neural populations in distinct regions of the non-primary auditory cortex; these neurons respond selectively to music and not speech or other environmental sounds^6,8,9^. Several experimental observations suggest that music-selectivity and an ability to process the basic features of music develop spontaneously, without special need for an explicit musical training^10^. For example, a recent neuroimaging study showed that music-selective neural populations exist in not only individuals who had explicit musical training but also in individuals who had almost no explicit musical training^11^. In addition, it was reported that even infants have an ability to perceive multiple acoustic features of music^12,13^, such as melody that is invariant to shifts in pitch level and tempo, similar to adults. One intuitive explanation is that passive exposure to life-long music may initialize the music-selective neural populations^11^, as hearing occurs even during pre-natal periods^14^. However, the basic machinery of music processing, such as harmonicity-based sound segregation, has been observed not only in Westerners but also in native Amazonians who had limited exposure to concurrent pitches in music^15^. These findings raise speculations on whether exposure to music is necessary for the development of music-selectivity and how the universality of music can arise in different cultures.

Recent modeling studies using artificial deep neural networks (DNNs) have provided insights into the principles underlying the development of the sensory functions in the brain^16–19^. In particular, it was suggested that a brain-like functional encoding of sensory inputs can arise as a by-product of optimization to process natural stimuli in DNNs. For example, responses of DNN models trained for classifying natural images were able to replicate visual cortical responses and could be exploited to control the response of real neurons beyond the naturally-occurring level^20–22^. Even high-level cognitive functions have been observed in networks trained to classify natural images, namely the Gestalt closure effect^23^ and the ability to estimate the number of visual items in a visual scene^24,25^. Furthermore, a DNN trained for classifying music genres and words was shown to replicate human auditory cortical responses^26^, implying that such task-optimization provides a plausible means for modeling the functions of the auditory cortex. Based on this, we investigated a scenario in which music-selectivity can arise as a by-product of adaptation to natural sound processing in neural circuits^27–30^, so that the statistical patterns of natural sounds constrain universals of music in our brain.

We initially tested whether a distinct representation of music can arise in a DNN trained for detecting natural sounds (including music) using the AudioSet dataset^31^. Previous work suggested that a DNN trained to classify music genres and word categories can explain the responses of the music-selective neural populations in the brain^26^. Thus, it was expected that DNNs can learn general features of music to distinguish them from diverse natural sound categories.

The dataset we used consists of 10 s real-world audio excerpts from YouTube videos that have been human-labeled with 527 categories of natural sounds (**Fig. 1A**, 17,902 training data and 17,585 test data with balanced numbers for each category to avoid overfitting for a specific class). The design of the network model (**Fig. 1B** and **Table S1**) is based on conventional convolutional neural networks^32^, which have been employed to successfully model both audio event detection^33^ and information processing of the human auditory cortex^26^. The network was trained to detect all audio categories in each 10 s excerpt (e.g., music, speech, dog barking, etc.). As a result, the network achieved reasonable performance in audio event detection as shown in **Fig. S1A**. After training, 17,585 test data was presented to the network and the responses of the units in the average pooling layer were used as feature vectors representing the data.

**Fig. 1.**
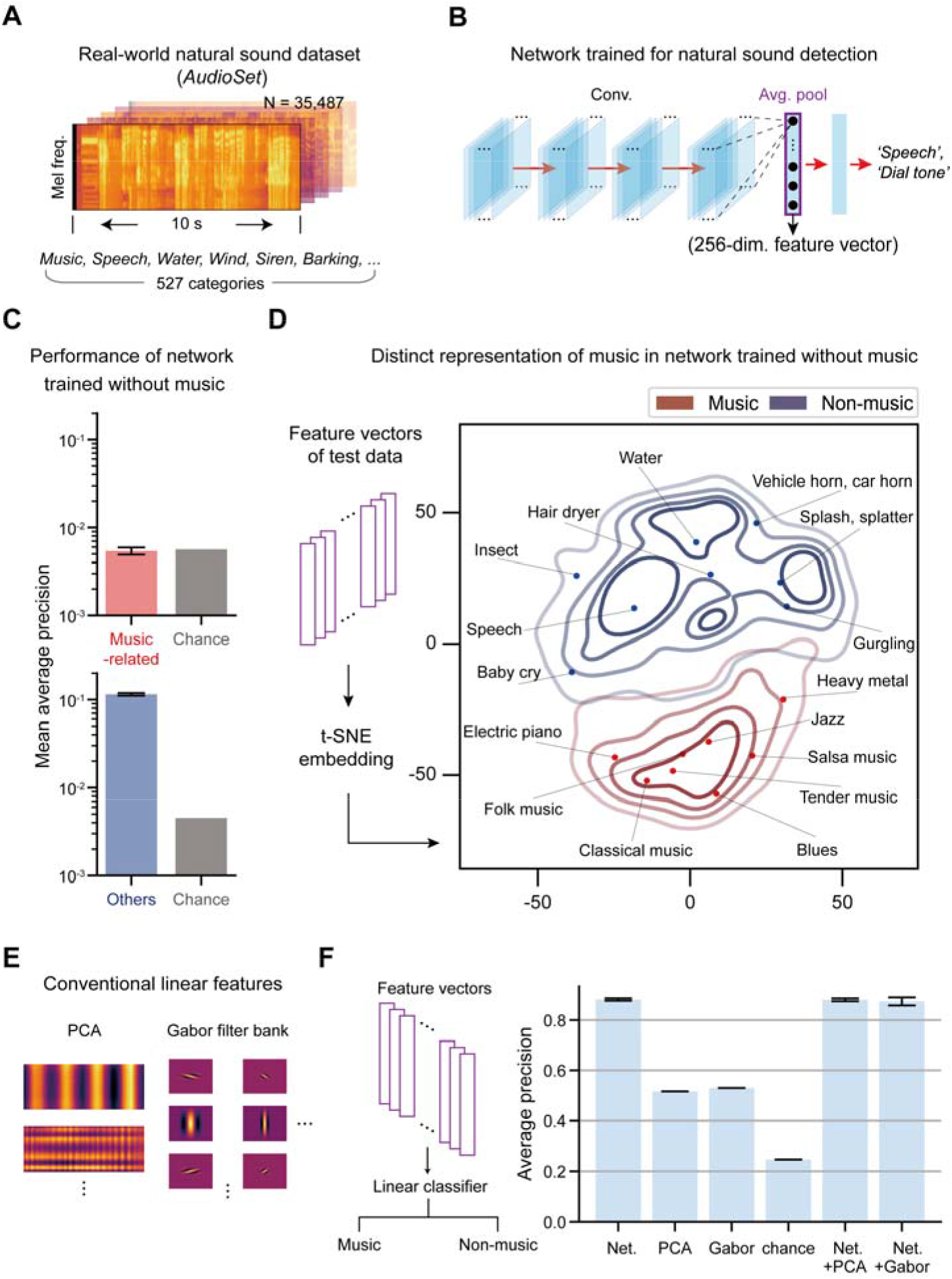
Distinct representation of music in deep neural networks trained for natural sound detection with and without music. (A) Example log-Mel spectrograms of the natural sound data in AudioSet^31^. (B) Architecture of the deep neural network used to detect the natural sound categories in the input data. The purple box indicates the average pooling layer. (C) Performance (mean average precision, mAP) of the network trained without music for music-related categories (top, red bars) and other categories (bottom, blue). (D) Density plot of the t-SNE embedding of feature vectors obtained from the network in C. The lines represent iso-proportion lines at 80%, 60%, 40%, and 20% levels. (E) Two conventional methods for linear feature extraction. Examples of principal components (left) and Gabor filters (right) are shown. (F) Binary classification of the data using a linear regression classifier. Error bars represent the standard deviation for different network initialization conditions in (C) and (F).

By analyzing the feature vectors of music and non-music data, we confirmed that the network trained with music has a unique representation for music, distinct from other sounds. We used t-distributed stochastic neighbor embedding (t-SNE) to visualize the 256-dimensional feature vectors in two dimensions, which ensures that data close in the original dimensions remain close in two dimensions^34^. The resulting t-SNE embedding shows that the distribution of music data is clustered in a distinct territory of the embedding space, clearly separated from non-music data (**Fig. S1B**). Such a result is expected; as music was included in the training data, the network can learn the features of music that distinguish music from other categories. Given this, one might expect that such a distinct representation of music would not appear if music were discarded from the training dataset.

However, further investigation showed that the distinct representation for music can still arise in a DNN trained without music. To test this, we discarded the data that contain any music-related categories from the training dataset and trained the network to detect other audio events except the music-related categories. As a result, the network was not able to detect music-related categories, but still achieved reasonable performance in other audio event detection (**Fig. 1C**).

Interestingly though, the distribution of music was still clustered in a distinct regime of the t-SNE embedding space, despite the network not being trained with music (**Fig. 1D**). We quantified such separation by calculating the segregation index (SI) between music and non-music in the t-SNE space. The SI of the network trained with natural sound excluding music was comparable to that of the network trained with natural sound including music (**Fig. S1B**), implying that training with music is not necessary for the distinct representation of music by the DNN.

Such observation raises a question on how such distinct representations emerge without training music. Based on previous notions^27–30^, we speculated that features important for processing music can spontaneously emerge as a by-product of learning natural sound processing in DNNs. To rule out other possibilities first, we tested two alternative scenarios: 1) music and non-music can be separated in the representation space of the log-Mel spectrogram using linear features, so that a nonlinear feature extraction process is not required, and 2) units in the network selectively respond to the trained categories but not to unseen categories, so that the distinct representation emerges without any music-related features in the network.

We first confirmed that the distinct representation did not appear when conventional linear models were used. To test this, feature vectors were obtained from data in the log-Mel spectrogram space by applying two conventional models for linear feature extraction: principal component analysis (PCA, **Fig. 1E** and **Fig. S2A**) and a spectro-temporal two-dimensional-Gabor filter bank (GBFB) model of auditory cortical response^35,36^ (**Fig. 1E** and **Fig. S2C**, Methods).

Next, we applied the t-SNE embedding method to the obtained vectors, as in **Fig. 1D**, and analyzed the distribution. The resulting embedding generated by the PCA and GBFB methods did not show a clear separation between music and non-music (**Figs. S2B and S2D**), while showing significantly lower SI values compared to the SI of networks trained without music (PCA: SI = 0.365, p= 0.031; GBFB: SI = 0.331, p = 0.031, Wilcoxon signed rank-sum test).

To further confirm this tendency while avoiding any distortion of data distribution that might arise from the dimension reduction process, we fitted a linear regression model to classify music and non-music in the training dataset by using their feature vectors as predictors and tested the classification performance using the test dataset (**Fig. 1F**). As a result, the network trained with natural sounds yielded significantly higher accuracy (mAP of network trained without music: 0.883 ± 0.005, chance level: 0.246) than PCA or GBFB (PCA: mAP = 0.515, p = 0.031; GBFB: mAP = 0.529, p = 0.031, Wilcoxon signed rank-sum test). Moreover, the classification accuracy was almost unchanged even when the linear features were used together with the features from the network (Net+PCA: mAP = 0.881 ± 0.006, Net+GBFB: mAP = 0.875 ± 0.016). These results suggest that conventional linear features cannot explain the distinct representation of music found in the embedding space.

Next, we tested whether the distinct representation is due to the specificity of the unit response to the trained categories^37,38^. It is possible that all features learned by the network are specifically fitted to the trained sound categories, so that the sounds of the trained categories would elicit a reliable response from the units while the sounds of unseen categories (including music) would not. To test this, we checked whether the average response of the units to music is significantly smaller than the non-music stimuli. Interestingly, the average response to music was stronger than the average response to non-music (**Fig. 2A**, p <10^−10^, Wilcoxon signed rank-sum test). This suggests that features optimized to detect natural sound can also be rich repertoires of music; i.e., the network may have learned features of music throughout the training process even though music was completely absent in the training data.

**Fig. 2.**
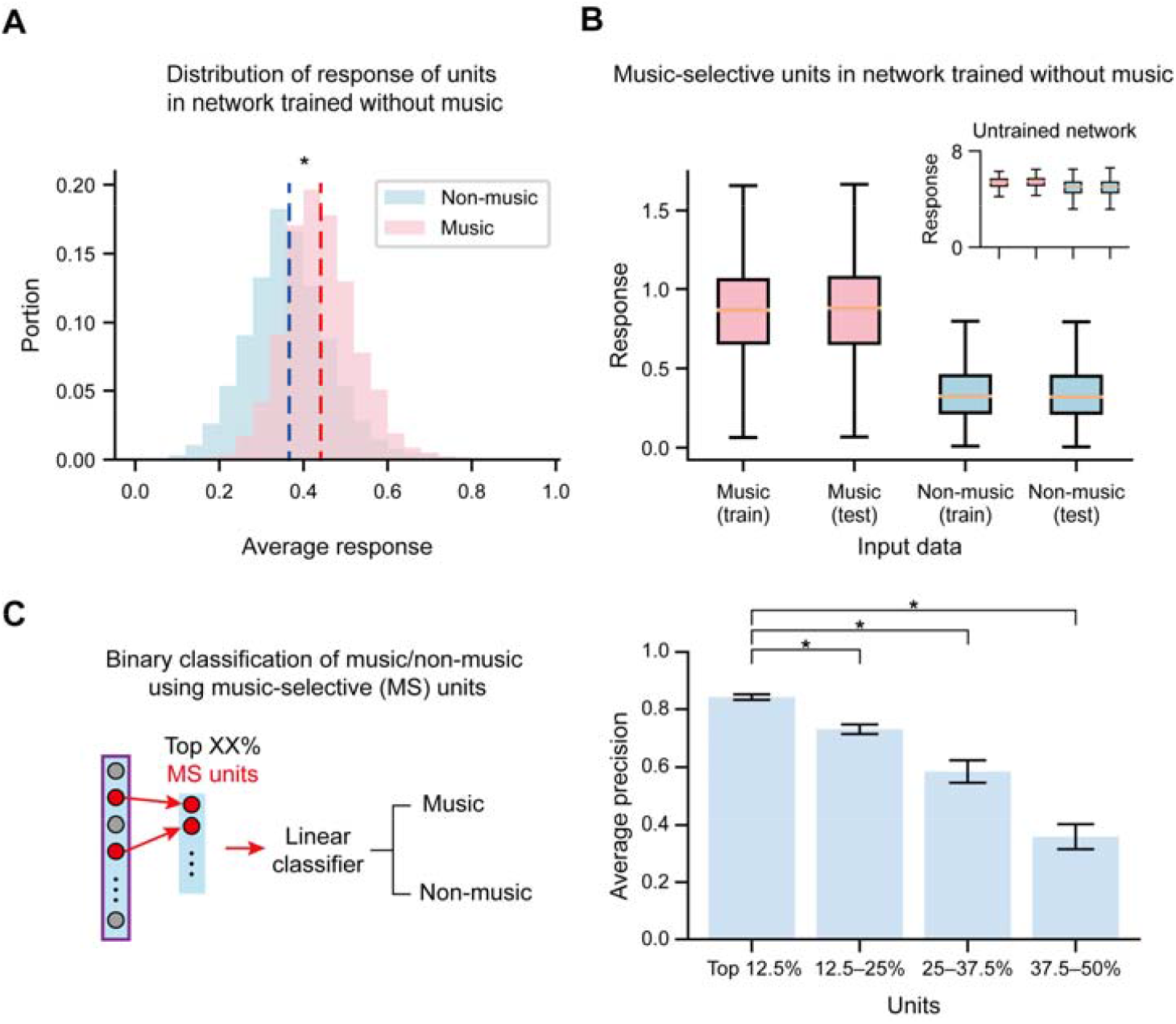
Selective response of units in the network to music. (A) Histograms of the response of the units averaged over music (red) and non-music (blue) stimuli in networks trained without music. The dashed lines represent the response averaged over all units. (B) Response of the music-selective units to music (red) and non-music stimuli. Inset: Response of the units in the untrained network with the top 12.5% MSI values to music and non-music stimuli. Error bars represent the standard deviation for various inputs. (C) Illustration of the binary classification of music and non-music using the response of the music-selective units (left), and the performance of the linear classifier (right). The asterisks indicate statistical significance (p<0.05) (from the top, p = 0.031, p = 0.031, p = 0.031, Wilcoxon signed rank-sum test). Error bars represent the standard deviation for different network initialization conditions.

Based on the above results, we investigated whether units in the network exhibit music-selective responses. We used two criteria to confirm this: 1) whether some units show a significantly stronger response to music than other sounds, and 2) whether those units encode the temporal structure of music in multiple timescales.

First, we confirmed that some units in the network respond selectively to music rather than other sounds. To evaluate this, we define and quantify the music-selectivity index (MSI) of each network unit as the difference between the average response to music and non-music in the training dataset normalized by their unpooled variance^39^ (i.e., *t*-statistics, Methods). The units with the top 12.5% MSI values (MSI = 51.0 ± 9.6) showed a 2.76 times stronger response to music than other sounds in the test dataset on average (**Fig. 2B**), and thus were considered as putative music-selective units. We confirmed that the response of these music-selective units can be exploited for the music classification task (**Fig. 2C**, accuracy: AP: 0.842 ± 0.010) using a linear classifier as in **Fig. 1F**. In contrast, using other units with intermediate MSI values showed significantly lower performance (top 37.5–50%, AP: 0.359 ± 0.044, p = 0.031, Wilcoxon signed rank-sum test), confirming that the music-selective units provide useful information for processing music.

Second, we found that the music-selective units in the network showed sensitivity to the temporal structure of music, replicating previously observed characteristics of tuned neural populations in the human auditory cortex^6,40,41^. While music is known to have distinct features in both long and short timescales^6,41^, it is possible that the putative music-selective units only encode specific features of music in a specific (especially short) timescale. To test this, we adopted the ‘sound quilting’ method^41^ (**Fig. 3A**, Methods), as follows: the original sound sources were divided into small segments (50–1,600 ms in octave range) and then reordered while considering smooth connections between segments. This shuffling method preserves the acoustic properties of the original sound on a short timescale but destroys it on a long timescale. It has been shown that music-selective neural populations in the human auditory cortex respond robustly when the segment size is large (e.g., 960 ms) so that most of the temporal structures are preserved, but the response is greatly reduced when the segment size is small (e.g., 30 ms) so that the temporal structure of the original sound is broken^41^. Similarly, after recording the response of the music-selective units to such sound quilts of music, we confirmed that their response is strongly correlated with the segment size (music quilt: r = 0.57, p = 0.00093). The response was mostly similar to the case of giving the original sound as an input in 800 ms segments, but greatly reduced when 50 ms segments were given (**Fig. 3B**, original: 0.743 ± 0.043; 800 ms: 0.751 ± 0.042; 50 ms: 0.569 ± 0.028; p_original-50 ms_ = 0.031, p_original-800 ms_ = 0.91, Wilcoxon signed rank-sum test).

**Fig. 3.**
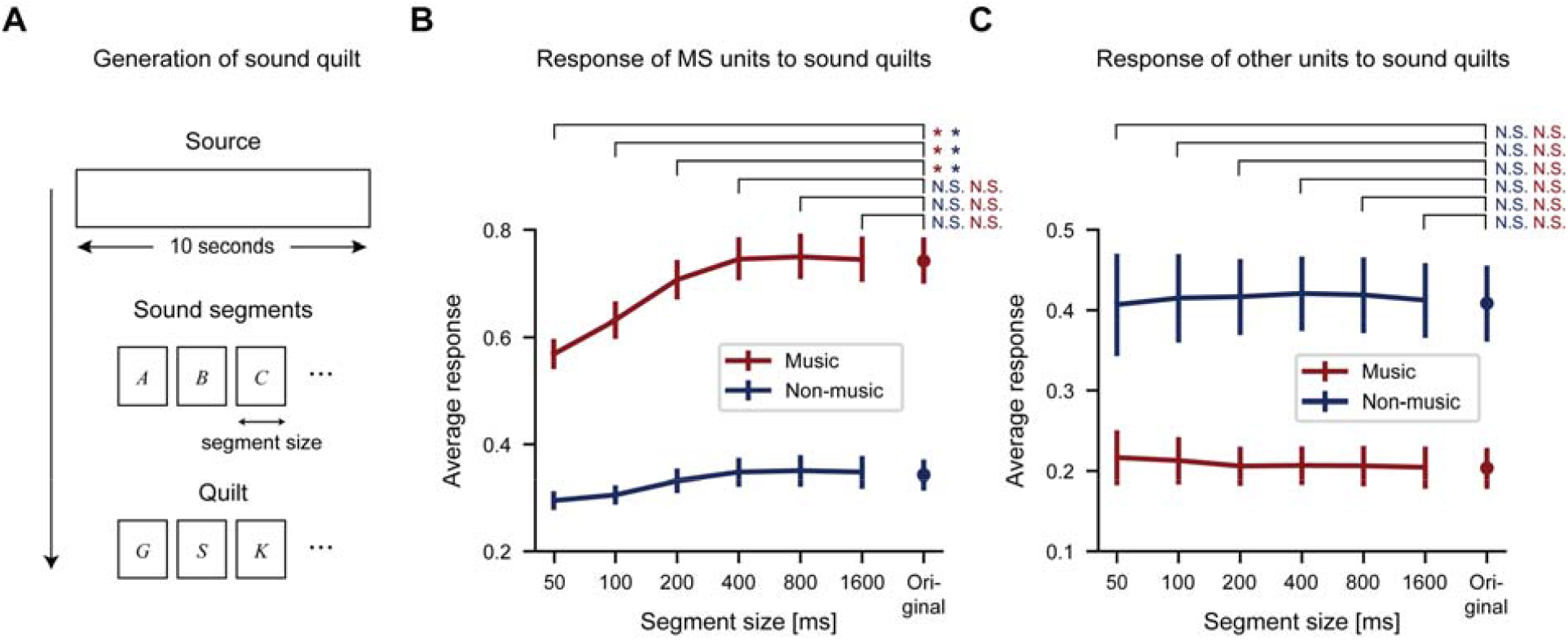
Encoding of the temporal structure of music by music-selective units in the network as in the human brain. (A) Schematic diagram of the generation of sound quilts. (B) Response of the music-selective units to sound quilts made of music (red) and non-music (blue). For the music quilts, from the top: p = 0.031, p = 0.031, p = 0.031, p = 0.69, p = 1.0, p = 0.91; for the non-music quilts, from the top: p = 0.031, p = 0.031, p = 0.031, p = 0.97, p = 1.0, p = 1.0, Wilcoxon signed rank-sum test. (C) Response of the other units to sound quilts made of music (red) and non-music (blue). For the music quilts, from the top: p = 0.91, p = 0.94, p = 0.84, p = 0.91, p = 0.91, p = 0.68; for the non-music quilts, from the top: p = 0.5, p = 0.84, p = 1.0, p = 1.0, p = 1.0, p = 0.91. The asterisks indicate statistical significance (p < 0.05). N.S.: non-significant (p > 0.05).

To test whether or not the effect is due to the quilting process itself, we provided quilts of music to the other non-music-selective units. In this condition, we confirmed that the average response remains constant even when the segment size changes (**Fig. 3C**). Furthermore, when quilted natural sound inputs were provided, the correlation between the response of the music-selective units and the segment length was weaker than when quilted music inputs were provided (**Fig. 3B**, non-music quilt: r = 0.45, p = 0.011), even though the significant correlation was observed for both types of inputs. Notably, all these characteristics of the network trained without music replicate those observed in the human brain^6,41^.

Then how does music-selectivity emerge in a network trained to detect natural sounds even without training music? In the following analysis, we found that music-selectivity can be a critical component to achieve generalization of natural sound in the network, and thus training to detect natural sound spontaneously generates music-selectivity.

Clues were found from the observation that the music-selectivity of the network gradually increases throughout the training process for natural sound detection. We measured both SI and task performance of networks over the course of training (**Fig. S3A**) and found that both SI and task performance monotonically increase and saturate at approximately 30 training epochs (**Fig. S3B**). Accordingly, we confirmed that SI and task performance are strongly correlated (**Fig. S3C**, from 0 to 50 epochs, r = 0.76, p = 5.2 × 10^−49^), implying that a network’s natural sound detection performance can be used to predict its music-selectivity.

Based on this, we hypothesized that music-selectivity can act as a functional basis for the generalization of natural sound, so that the emergence of music-selectivity may directly stem from the ability to process natural sounds. To test this, we investigated whether music-selectivity emerges when the network cannot generalize natural sounds (**Fig. 4A**). To hinder the generalization, the labels of the training data were randomized to remove any systematic association between the sound sources and their labels, following a previous work^42^. Even in this case, the network achieved high training accuracy (training AP > 0.95) by memorizing all the randomized labels in the training data, but showed a test accuracy at the chance level as expected.

**Fig. 4.**
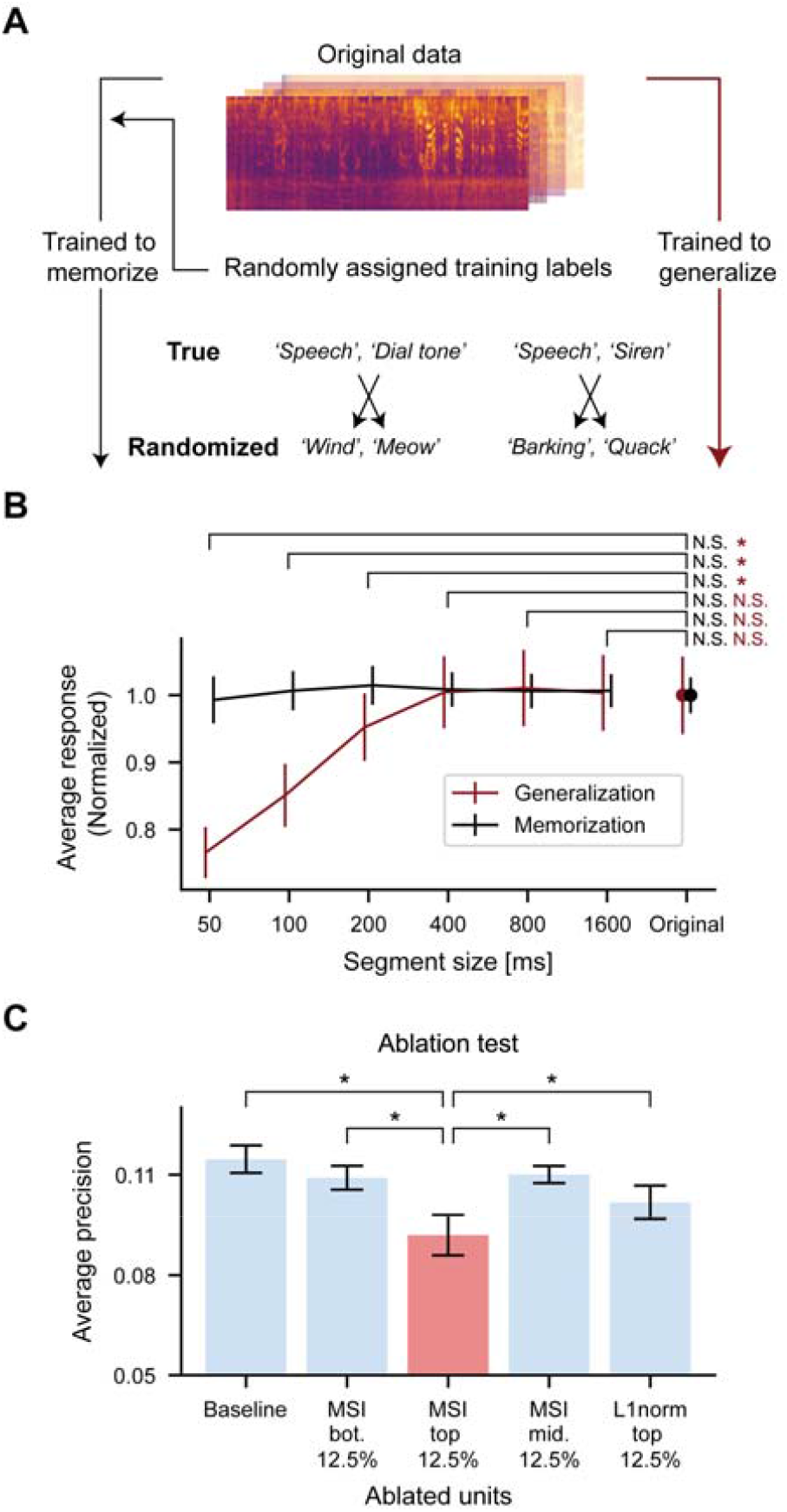
Music-selectivity as a generalization of natural sounds. (A) Illustration of network training to memorize the data by randomizing the labels. (B) Response of the units with the top 12.5% MSI values to music quilts in the networks trained with randomized labels (black, memorization) compared to that of the network in Fig. 3B (red, generalization). To normalize the two conditions, each response was divided by the average response to the original sound from each network. For memorization, from the top: p = 0.41, p = 0.69, p = 0.97, p = 1.0, p = 1.0, p = 0.94, Wilcoxon signed rank-sum test. N.S.: non-significant (p > 0.05). Error bars represent the standard deviation for different network initialization conditions. (C) Performance of the network after the ablation of specific units (red: ablation of music-selective units). From the top, from the left, p = 0.031, p = 0.031, p = 0.031, p = 0.031, Wilcoxon signed rank-sum test. The asterisks indicate statistical significance (p < 0.05).

We confirmed that the process of generalization is indeed critical for the emergence of music-selectivity in the network. For the network trained to memorize the randomized labels, the distributions of music and non-music were less distinct in the t-SNE embedding space compared to the network trained to generalize (**Fig. S4**, trained to memorize: SI = 0.587 ± 0.045, p = 0.0090, Wilcoxon rank-sum test), although some degree of separation was still observed. More importantly, units in the network trained to memorize did not encode the temporal structure of music. To test this, we analyzed the response of the units with the top 12.5% MSI values in the network trained to memorize using sound quilts of music as in **Fig. 3B**. We found that even if the segment size of the sound quilt changed, the response of the units remained mostly constant, unlike the music-selective units in the network trained to generalize natural sounds (**Fig. 4B**). This supports our hypothesis that music-selectivity is based on the process of generalization of natural sounds.

To further investigate the functional association, we performed an ablation test (**Fig. 4C**), in which the response of the music-selective units is silenced and then the sound event detection performance of the network is evaluated. If the music-selective units provide critical information for the generalization of natural sound, removing their inputs would greatly reduce the performance of the network. Indeed, we found that ablation of the music-selective units significantly deteriorates the performance of the network (**Fig. 4C**, red: top 12.5% music-selective units, performance drop = 19.7%, p_MSI top 12.5%-Baseline_ = 0.031, Wilcoxon signed rank-sum test).

This effect was much weaker when the same number of units with intermediate/bottom MSI values were silenced (intermediate: p = 0.031, bottom: p = 0.031). Furthermore, the performance drop was even greater than that of ablating the units showing strong responses to inputs on average (top 12.5% L1 norm, performance drop = 8.0%, p_MSI top 12.5%-L1norm top 12.5%_ = 0.031, Wilcoxon signed rank-sum test). This suggests that music and other natural sounds share key features, and thus music-selective units can play a functionally important role not only in music processing but also in natural sound detection.

## Discussions

What is the origin of music? Here, we put forward the notion that neural circuits for processing the basic elements of music can develop spontaneously as a by-product of adaptation for natural sound processing. In the DNN trained for natural sound detection in this work, music was distinctly represented even when music was not included in the training data. Such distinction cannot be explained by conventional linear features, but rather arises from the response of the music-selective units in the feature extraction layer. The music-selectivity was also sensitive to the temporal structure of music, replicating all of the observed characteristics of the music-selective neural populations in the brain. Further investigation suggested that music-selectivity can work as a functional basis for the generalization of natural sound, revealing how it can emerge without learning music. All together, these results support the notion that a universal template of music can arise from evolutionary pressure to process natural sound.

Our model provides a simple explanation about why a DNN trained to classify musical genres replicated the response characteristics of the human auditory cortex^26^, although it is unlikely that the human auditory system itself has been optimized to process music. This is because training with music would result in learning general features for natural sound processing, as music and natural sound processing share a common functional basis. The existence of a basic ability to perceive music in multiple non-human species is also explained by the model. Our analysis showed that music-selectivity lies on the continuum of learning natural sound processing.

If the mechanism also works in the brain, such ability would appear in a variety of species adapted to natural sound processing, but to varying degrees. Consistent with this idea, the processing of basic elements of music has been observed in multiple non-human species: octave generalization in rhesus monkeys^45^, the relative pitch perception of two-tone sequences in ferrets^46^, and a pitch perception of marmoset monkeys similar to that of humans^47^.

Neurophysiological observations that neurons in the primate auditory cortex selectively respond to pitch^48^ or harmonicity^49^ were also reported, further supporting the notion. A further question is whether phylogenetic lineage would reflect the ability to process the basic elements of music, as our model predicts that music-selectivity is correlated with the ability to process natural sounds.

Our results also provide insights into the workings of audio processing in DNNs. Recent works showed that the class selectivity of DNN units is a poor predictor of the importance of the units and can even impair generalization performance^51,52^, possibly because it can induce overfitting to a specific class. On the other hand, we found that music-selective units are important for the natural sound detection task, and a good predictor of DNN performance. One possible explanation is that the music-selective units have universal features for the generalization of other natural sounds rather than specific features for specific classes, and thus removing them greatly hinders the performance of the DNN. Thus, these results also support the notion that the general features of natural sounds learned by DNNs are key features that make up music.

In summary, we demonstrated that music-selectivity can spontaneously arise in a DNN trained with real-world natural sounds without music, and that the music-selectivity provides a functional basis for the generalization of natural sound processing. By replicating the key characteristics of the music-selective neural populations in the brain, our results encourage the possibility that a similar mechanism could occur in the biological brain, as suggested for visual^22– 24^ and navigational^53^ functions using task-optimized DNNs. Our findings support the notion that ecological adaptation may initiate various functional tunings in the brain, providing insight into how the universality of music and other innate cognitive functions arises.

## Materials and Methods

All simulations were done in Python using the PyTorch and TorchAudio framework.

### Neural network model

Our simulations were performed with conventional convolutional neural networks for audio processing. At the input layer, the original sound waveform (sampling rate = 22,050 Hz) was transformed into a log-Mel spectrogram (64 mel-filter banks in the frequency range of 0 Hz to 8,000 Hz, window length: 25 ms, hop length: 12.5 ms). Next, four convolutional layers followed by a batch-normalization layer and a max-pooling layer (with ReLU activation and a dropout rate of 0.2) extracted the features of the input data. The global average pooling layer calculated the average activation of each feature map of the final convolutional layer. These feature values were passed to two successive fully connected layers, and then a sigmoid function was applied to generate the final output of the network. The detailed hyperparameters are given in **Table S1**.

### Stimulus dataset

The dataset we used is the AudioSet dataset^31^, a collection of human-labeled (multi-label) 10 s clips taken from YouTube videos. We used a balanced dataset (17,902 training data and 17,585 test data from distinct videos) consisting of 527 hierarchically organized audio event categories (e.g., ‘classical music’ under ‘music’). Music-related categories were defined as all classes under the music hierarchy. Each excerpt in the dataset is intrinsically multi-labeled as different sounds generally co-occur in a natural environment, but a sufficient number of data was selected to contain only music-related categories (3,620 in the training set and 4,033 in the test set) and no music-related categories (11,087 in the training set and 10,616 in the test set). To test for the distinct representation of music, the data were reclassified into music, non-music, and mixed sound, and then mixed sounds were excluded in the analysis of music-selectivity. This was required because some data that contained music-related categories can also contain other audio categories (e.g., music + barking).

### Network training

We trained the network to detect all sound categories in each 10 s clip (multi-label detection task). To that aim, the loss function of the network was chosen as the binary cross-entropy between the target (y) and the output (x), which is defined as

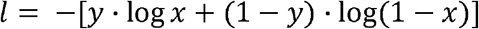

for each category. For optimizing this loss function, we employed the AdamW optimizer with weight decay = 0.01^54^. Each network was trained for 100 epochs (200 epochs for the randomized labels) with a batch size of 32 and the One Cycle learning rate (LR) method^55^. The One Cycle LR is an LR scheduling method for faster training and preventing the network from overfitting during the training process. This method linearly anneals the LR from the initial LR 4 × 10^−5^ to the maximum LR 0.001 for 30 epochs and then from the maximum LR to the minimum LR 4 × 10^−9^ for the remaining epochs. For every training condition, simulations were run for five different random seeds of the network. The network parameters used in the analysis were determined from the epoch that achieved the highest average precision over the training epochs with 10% of the training data used as a validation set.

### Analysis of the responses of the network units

The responses of the network units in the average pooling layer were analyzed as feature vectors (256 dimensions) representing the data. After t-SNE embedding (perplexity = 30) of the feature vectors, we measured the SI to quantify the separation between the probability distribution of music and non-music, which is defined as

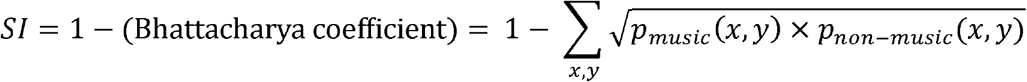

where *p* represents the probability distribution of music and non-music in t-SNE embedding space.

Following a previous experimental study^39^, the music-selectivity index of each unit was defined as

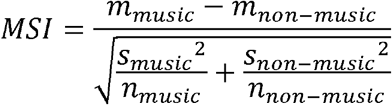

where *m* is the average response of a unit to music and non-music stimulus, *s* is the standard deviation, and *n* is the number of each type of data.

### Extraction of linear features using conventional approaches

The linear features of the log-Mel spectrogram of the natural sound data were extracted by using principal component analysis (PCA) and the spectro-temporal two-dimensional-Gabor filter bank (GBFB) model following previous works^35,36^. In the PCA case, feature vectors were obtained from the top 256 principal components (total explained variance: 0.965). In the case of the GBFB model, a set of Gabor filters were designed to detect specific spectro-temporal modulation patterns, which are defined as

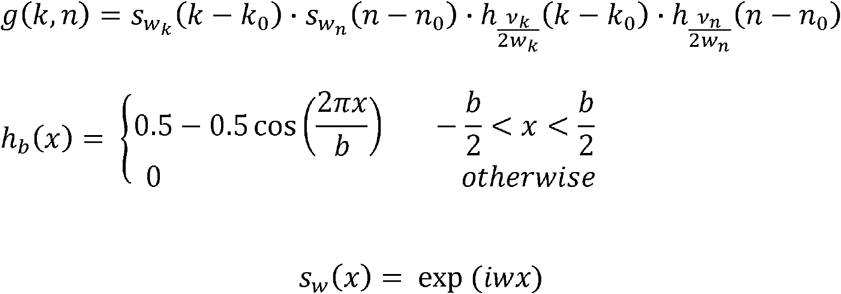

where *k* and *n* represent the channel and time variables (center: *k*_*0*_ and *n*_*0*_), *w*_*k*_ is the spectral modulation frequency, *w*_*n*_ is the temporal modulation frequency, and *v* is the number of semi-cycles under the envelope. The distribution of the modulation frequencies was designed to limit the correlation between filters as follows,

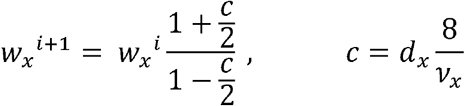

Here, we used *d*_*k*_ = 0.1, *d*_*n*_ = 0.045, *v*_*k*_ = *v*_*n*_ = 3.5, with *w*_*k, max*_ = *w*_*n, max*_ = π/4, resulting in 15 spectral modulation frequencies, 18 temporal modulation frequencies, and 263 independent Gabor filters (15×18–7). Next, a log-Mel spectrogram was convolved with each Gabor filter and then averaged to generate the 263-dimensional feature vector representing the data. Nonetheless, our investigation showed that the specific choice of the parameters does not change the results significantly.

### Generation of sound quilts

Sound quilts were created according to the algorithm proposed in a previous work^41^. First, the original sound sources were divided into small segments of equal size (50–1,600 ms in octave range). Next, these segments were reordered while minimizing the difference between the segment-to-segment change in log-Mel spectrogram of the original sound and that of the shuffled sound. Finally, we concatenated these segments while minimizing the boundary artifacts by matching the relative phase between segments at the junction ^41^.

### Ablation test

In the ablation test, the units in the network were grouped based on MSI value: top 12.5% units (MS units, N = 16), middle 43.75–56.25% units, and bottom 12.5% units. In addition, we grouped the units that showed a strong average response to the test data (top 12.5% L1 norm). The response of the units in each group was set to zero to investigate their contribution to natural sound processing.

### Statistical analysis

All statistical variables, including the sample sizes, exact *P* values, and statistical methods, are indicated in the corresponding texts or figure legends.

### Data and code availability

The data and codes that support the findings of this study are available at https://github.com/kgspiano/Music

## Supplementary Materials

Fig. S1. Distinct representation of music in deep neural networks trained for natural sound detection with music.

Fig. S2. T-SNE embedding of the feature vectors obtained by linear methods. Fig. S3. Correlation of music-selectivity and network performance.

Fig. S4. T-SNE embedding of the feature vectors of the network trained to memorize natural sounds with randomized labels.

Table S1. Summary of the network architecture.

## Figures and Tables

**Fig. S1.**
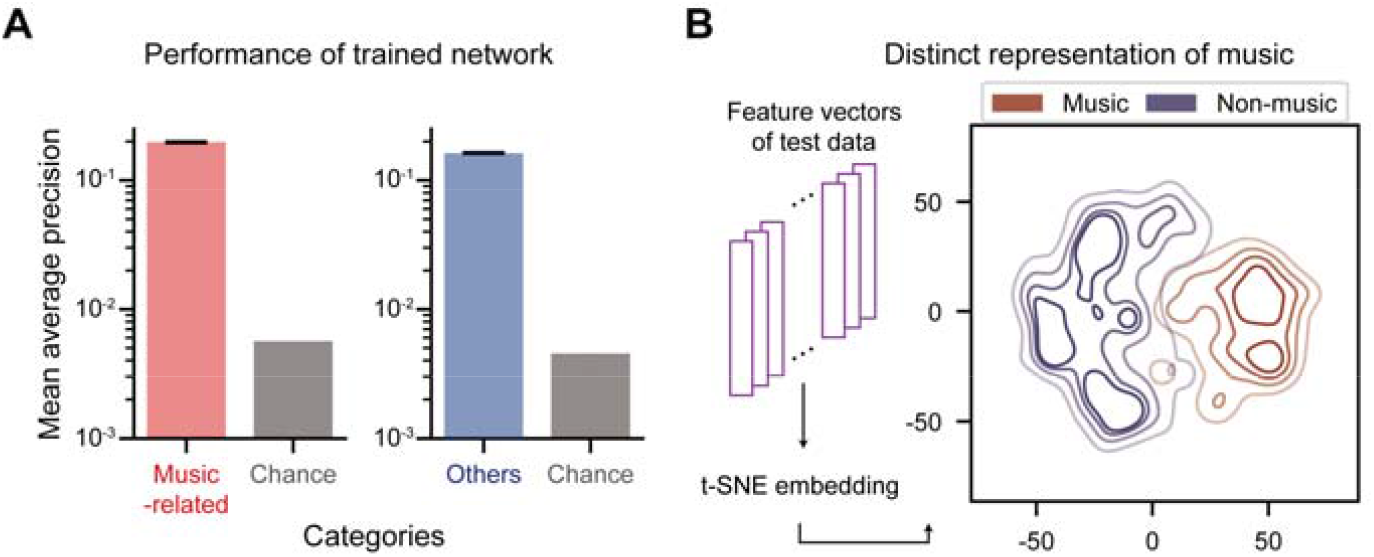
Distinct representation of music in deep neural networks trained for natural sound detection with music. (**A**) Performance of the trained network for music-related categories (left, red bars) and other categories (right, blue). (**B**) Density plot of the t-SNE embedding of feature vectors obtained from the trained network. The lines represent iso-proportion lines at 80%, 60%, 40%, and 20% levels.

**Fig. S2.**
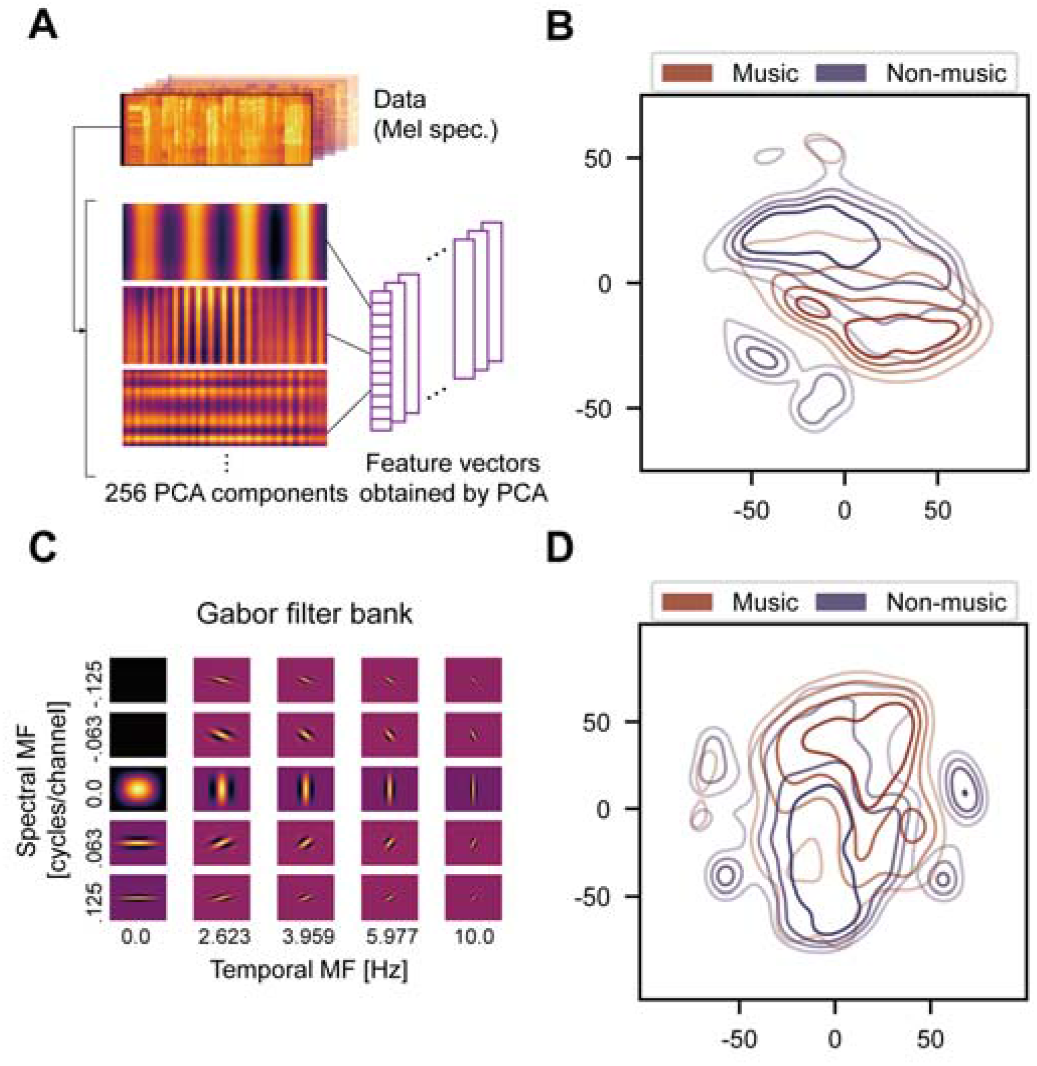
T-SNE embedding of the feature vectors obtained by linear methods. (A) Example PCA components obtained from the data. (B) Density plot of the t-SNE embedding of feature vectors obtained from PCA. The lines represent iso-proportion lines at 80%, 60%, 40%, 20% levels. (C) Example spectro-temporal Gabor filters. MF: modulation frequency. (D) Density plot of the t-SNE embedding of feature vectors obtained from Gabor filters.

**Fig. S3.**
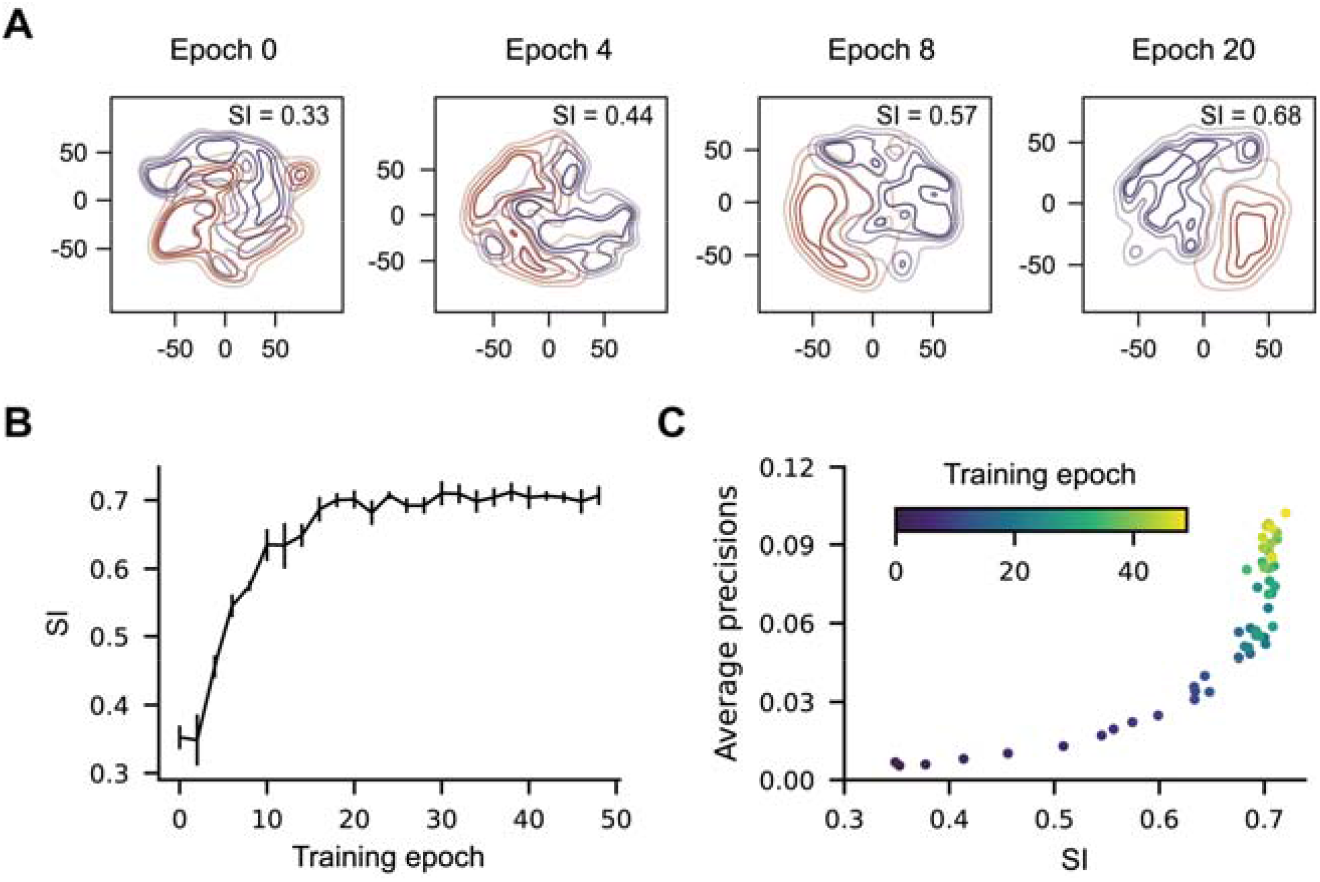
Correlation of music-selectivity and network performanc. Density plots of the t-SNE embeddings of music (red) and non-music (blue) over the training epochs. SI vs. training epoch, and (C) average precision vs. SI, showing that the segregation index and task performance are strongly correlated. Error bars represent the standard deviation for different network initialization conditions.

**Fig. S4.**
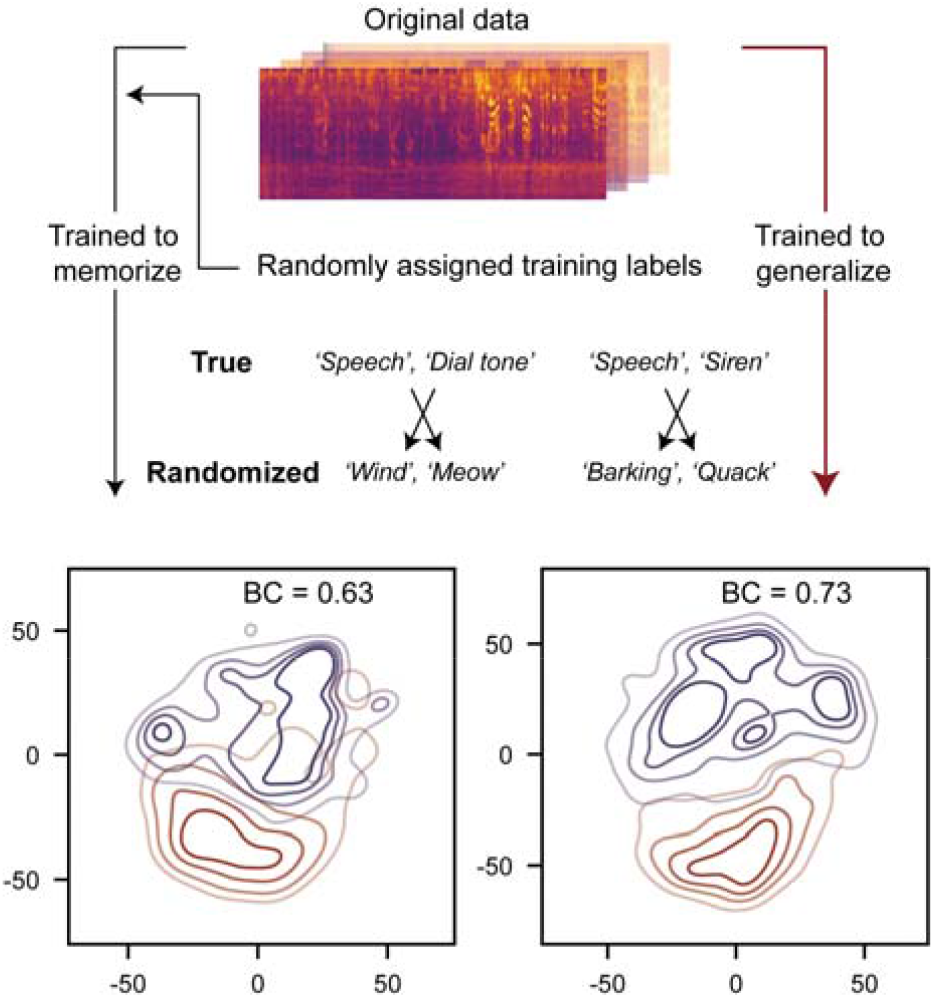
T-SNE embedding of the feature vectors of the network trained to memorize natural sounds with randomized labels. (top) Illustration of network training to memorize the data by randomizing the labels. (left) Density plot of the t-SNE embedding of the feature vectors obtained from the network trained with randomized labels and (right) with the original labels.

**Table S1.** Summary of the network architecture. The network consists of four convolutional layers for feature extraction (Conv1 – Conv4) and two fully connected layers for natural sound detection (FC1 – FC2). We note that the specific choice of hyperparameters does not significantly change the results in the main text.

| Layer | Type | Output Shape | Kernels | Activations |
| --- | --- | --- | --- | --- |
| Input | Log-Mel spectrogram input | 64 × 802 × 1<br>(height × width × channel) |  |  |
| Conv1 | Convolution | 30 × 200 × 32 | Size: 5 × 5 × 1 × 32<br>Stride: 2 × 4 | Batch normalization and ReLU |
| Pool1 | Max pooling | 15 × 100 × 32 | Size = 2 × 2<br>Stride = 2 | Dropout (p = 0.2) |
| Conv2 | Convolution | 11 × 96 × 64 | Size: 5 × 5 × 32 × 64<br>Stride: 1 | Batch normalization and ReLU |
| Pool2 | Max pooling | 10 × 95 × 64 | Size = 2 × 2<br>Stride = 1 |  |
| Conv3 | Convolution | 6 × 91 × 128 | Size: 5 × 5 × 64 × 128<br>Stride = 1 | Batch normalization and ReLU |
| Pool3 | Max pooling | 5 × 90 × 128 | Size: 2 × 2<br>Stride: 1 | Dropout |
| Conv4 | Convolution | 1 × 86 × 256 | Size: 5 × 5 × 192 × 256<br>Stride: 1 | Batch normalization, ReLU, and dropout |
| AvgPool1 | Global average pooling | 1 × 1 × 256 |  |  |
| FC1 | Fully Connected | 256 | Weights: 256 × 256<br>Bias: 256 × 1 | ReLU and dropout |
| FC2 (Output) | Classification Output | 527 | Weights: 527 × 256<br>Bias: 527 × 1 | Sigmoid |

## Notes

### Competing Interest Statement

The authors have declared no competing interest.

### Summary of Updates

Revised figure. Errata.

